# Saturated cardiolipins are potent disruptors of inner mitochondrial membrane structure and function

**DOI:** 10.64898/2026.04.14.718012

**Authors:** Kailash Venkatraman, Daniel Milshteyn, Carolina Sarto, Elida Kocharian, Cailyn M. Sakurai, Aaron M. Armando, Adrian M. Wong, Edward A. Dennis, Xi Fang, Christopher T. Lee, Itay Budin

## Abstract

Cardiolipin (CL) is a four-acyl chained, mitochondrial-specific phospholipid crucial for maintenance of inner mitochondrial membrane (IMM) structure and function. In healthy tissues, CL acyl chains are highly unsaturated and maintained by a conserved remodeling pathway. However, dysregulation of CL acyl chain composition can arise from mutations in the CL transacylase, Tafazzin (TAZ), resulting in Barth syndrome (BTHS), where patients exhibit heightened mitochondrial dysfunction. Cells lacking TAZ accumulate three-acyl chained monolysocardiolipin (MLCL) as well as CL species with saturated acyl chains (CL_sat_). While the presence of MLCL destabilizes electron transport chain (ETC) complexes and IMM-shaping proteins, the contributions of CL_sat_ to mitochondrial dysfunction have not been elucidated. Here, we find that treatment of TAZ knockout cells with exogenous saturated fatty acids causes accumulation of CL_sat_ and loss of IMM structure despite only minimal changes in MLCL composition. Imaging of cells with elevated CL_sat_ showed reduced fluidity of the inner membrane. Biophysical measurements and molecular dynamics analyses showed that di-saturated (C16:0 18:1)_2_ CL species order and rigidify membranes, while also losing the intrinsic lipid curvature characteristic of tetra-unsaturated CL. These results implicate CL_sat_ as a potential driver of mitochondrial dysfunction and an additional therapeutic target in mitigating BTHS pathology.

## Introduction

Cardiolipin (CL) is an anionic phospholipid (PL) synthesized and localized in the inner mitochondrial membrane (IMM), where it accounts for ∼20% of total mitochondrial PL content (1–4). Unlike all other PLs, CL exhibits a dimeric structure consisting of two headgroup phosphates and four acyl chains (5). Due to the difference in cross-sectional area between its headgroup and acyl chains, CL forms an inverted conical structure with a negative spontaneous curvature (3, 4). *In vitro* and *in silico* studies have demonstrated that CL clusters into regions of high curvature (4, 6, 7). In cells, CL is required for electron transport chain (ETC) stability (8, 9), supercomplex assembly (9), cristae formation (10, 11), and mitochondrial fission-fusion dynamics (12, 13). In energetic tissues such as the heart, CL is typically tetra-unsaturated with four linoleic (C18:2) acyl chains (14, 15). Unsaturated CL acyl chains are maintained by a conserved remodeling pathway involving repeated cycles of deacylation and transacylation (16). In mammals, immature CL synthesized *de novo* is deacylated to three-chained monolysocardiolipin (MLCL) by phospholipase A_2_ enzymes (17, 18) or by the alpha/beta hydrolase domain containing protein 18 (ABHD18) (19, 20). The transacylase TAZ mediates reacylation of MLCL to mature, unsaturated CL by transferring unsaturated acyl chains from other mitochondrial PLs (21, 22) (Figure 1A).

**Figure 1:**
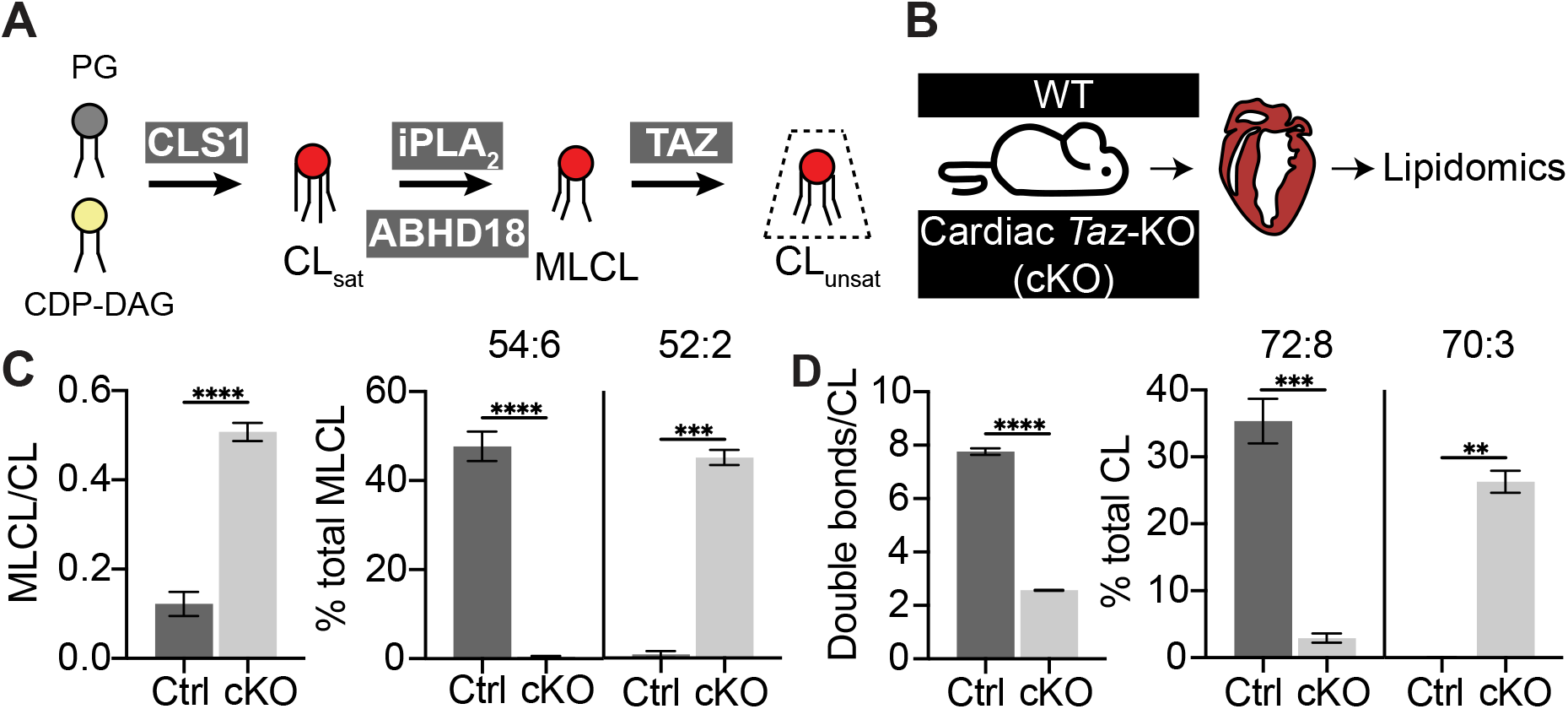
Cardiac-specific knockouts of Tafazzin alter the acyl chain profile of cardiolipin. (A) A conserved remodeling process ensures homogeneously unsaturated acyl chains on CL molecules. Unsaturated CL molecules exhibit a large magnitude of intrinsic curvature and are therefore depicted as having an inverted conical structure. The curvatures of MLCL and CL_sat_ species have not been experimentally determined. (B) Schematic representation of lipidomics analysis of cardiomyocyte-specific *Taz* knockouts (*Taz-*cKO) from whole mouse hearts. Lipidomics data was originally collected and described in (41). (C) *Taz-*cKO hearts exhibit accumulation of MLCL and an increase in MLCL with more saturated acyl chains. Lipidomic analysis was performed on n=3-4 mice per group. ****P<0.0001 and ***P=0.0001 by an unpaired two-tailed t-test. Error bars indicate standard deviation (SD). (D) *Taz-*cKO hearts lose unsaturated CL species and accumulate CL_sat_ species. ****P<0.0001, ***P=0.0002 and **P=0.0013 by an unpaired two-tailed t-test. Lipidomic analysis was performed on n=3-4 mice per group. Error bars indicate SD.

Dysregulation of CL acyl chain composition has been observed in numerous disorders (23), including Barth syndrome (BTHS). BTHS is a well characterized X-linked disorder that occurs upon mutations in TAZ, where patients present with dilated cardiomyopathy, skeletal myopathy as well as neutropenia (24–27). BTHS patients’ exhibit four major alterations in CL composition: 1) MLCL accumulation, 2) reductions in total CL, 3) increases in saturated CL species and 4) reduced levels of unsaturated CL (28–30). The accumulation of MLCL is the best characterized of the alterations in CL composition in BTHS. MLCL has been demonstrated to have weaker associations with mitochondrial proteins compared to unsaturated CL (31–33). Molecular dynamics (MD) studies on multiple scales show that MLCL exhibits reduced localization to regions of high curvature (34, 35), and reduced membrane deformation capabilities, compared to unsaturated CL species (36). In addition, MLCL has been shown to form altered peroxidase complexes with cytochrome c, contributing to increased oxidation of CL and other polyunsaturated fatty acids (37). Cellular models of BTHS, in which TAZ is disrupted or knocked out (TAZ-KO), show reduced membrane potential (38), supercomplex assembly (39, 40), and overall levels of ETC proteins (39, 40). However, they often lack some of the phenotypes observed in patients and animal models, like loss of highly-folded cristae structures in the IMM of BTHS patient lymphoblasts and bone marrow neutrophils (26, 41, 42).

Given the extensive characterization of the defective physicochemical properties of MLCL, inhibition of its formation by lipases that act on CL has been a primary avenue for seeking treatments for BTHS. The phospholipase A_2_ inhibitor bromoenol lactone (BEL) has been shown to restore the MLCL:CL ratio in some cellular models of BTHS, but it did not restore respiratory capacity (17, 43), suggesting that MLCL may not be the only molecular driver of mitochondrial dysfunction in BTHS. Notably, BEL-treated cells accumulate saturated CL species (CL_sat_) (18), which are products of *de novo* synthesis by cardiolipin synthase (CRLS) when remodeling is lost. A recent study developed an inhibitor for ABHD18 and showed a rescue of MLCL:CL ratio, but still observed accumulation of CL_sat_ due to loss of remodeling (20). In yeast, accumulation of CL_sat_ due to loss of the CL phospholipase A_2_ Cld1p causes loss of respiratory capacity and IMM structure when cells produce higher levels of saturated fatty acids due to oxygen limitation (44). These studies suggest that an underlying function of the remodeling pathway would be to buffer CL’s acyl chain composition (CL_sat_ levels) against changes in the cellular fatty acid pool, which can vary from diet and metabolic changes (45).

The properties of CL_sat_ and its functions have been challenging to disentangle from those of MLCL. Here we show that fatty acid supplementation in a BTHS cellular model primarily acts to alter levels of CL_sat_, likely by acting on CL precursor phospholipids. We observed that TAZ-KO C2C12 cells fed with palmitic (Palm), but not oleic acid (OA), accumulate CL_sat_ independently of changes in MLCL abundance, which corresponds to a loss of IMM structure, properties, and respiratory capacity. Motivated by these findings, we describe the biophysical properties of different CL species, which can be dramatically altered by the incorporation of only two saturated acyl chains. Our findings reveal that CL_sat_ species, in addition to MLCL, can drive alterations to membrane physiochemistry and mitochondrial function that could be relevant to pathologies related to CL metabolism.

## Results

Accumulation of both MLCL and CL_sat_ species are hallmarks of BTHS patient lipidomes and mouse models of BTHS. Cardiomyocyte-specific knockouts of Tafazzin (*Taz*-cKO) mice mimic the cardiac defects exhibited in BTHS patients (Figure 1B) (41). Isolated hearts from *Taz*-cKO mice exhibit major increases in MLCL, as well as accumulation of more saturated MLCL species such as 52:2 (Figure 1C) (46). Notably, *Taz*-cKO mice also exhibit major increases in the saturation of CL (8 double bonds to 2 per CL) species, with saturated 70:3 species as the most abundant (Figure 1D). Tetra-unsaturated CLs (i.e 72:8) were also significantly reduced compared to controls (Figure 1D). These changes were similar to those observed in studies using BTHS patient-derived lymphoblasts, where major increases in CL_sat_, with (C16:0 18:1)_2_ CL (POCL) compared to control cell lines were measured (47). CL saturation in *Taz*-cKO mice was similar to that of the CL-precursor PG, while CL was markedly more unsaturated in control animals (Figure S1A). This may represent the accumulation of de novo synthesized CL, whose acyl chain composition is similar to that of its substrates.

Based on the analysis of *Taz*-cKO mice, we hypothesized that levels of CL_sat_ would be sensitive to the cellular fatty acid pool in cells lacking Tafazzin activity, as that would determine de novo CL composition. Using a C2C12 myoblast model (38), we tested how WT and TAZ-KO cells would differentially alter CL acyl chain composition in response to mild, 100 μM, supplementation of Palm or OA (Figure 2A). This concentration of Palm-supplementation does not alter the morphology of mitochondria or other organelles (48), despite reducing the number of double bonds per PL acyl chain in the total PL pool (Figure 2A). Palm supplementation altered the composition of CL: WT cells exhibited a modest increase in CL_sat_ when compared to OA, and a 15% increase in POCL abundance (Figure 2B-C). This effect was magnified in Palm-treated TAZ-KO cells, with a decrease in CL unsaturation (4DB/CL to 3DB/CL) and a ∼60% increase in POCL abundance, the most abundant CL species in TAZ-KO cells, compared to OA-treated cells (Figure 2C). In addition, Palm-treated WT and TAZ-KO cells both exhibited reductions in tetra-unsaturated (C18:1)_4_, (C16:1)(C18:1)_3_, and mono-saturated (C16:0)(C18:1)_3_ CL species (Figure 2C). Notably, CL saturation in TAZ-KO cells treated with Palm was similar to the saturation of the PG precursor pool (Figure S1B). On the other hand, WT cells treated with Palm exhibited an increase in PG saturation compared to OA-treated cells but still retained highly unsaturated CL, while OA-treated TAZ-KO cells showed high CL and PG unsaturation (Figure S1B). Indicating that defects to CL remodeling may sensitize CL composition to fatty acid supplementation by acting on CL precursors. Notably, TAZ-KO cells in both treatment conditions exhibited increases to PG levels, consistent with observations in *Taz-*cKO mouse hearts (Figure S1C-D). TAZ-KO cells fed with Palm still accumulated MLCL (Figure 2D) but did not exhibit a significant increase in MLCL abundance compared to OA-treated TAZ-KO cells (Figure 2D). Palm treatment in TAZ-KO cells also increased the saturation of MLCL, with (C16:0)(18:1)_2_ species accumulating instead of (C18:1)_3_ (Figure 2E). Thus, these treatments illustrate a modality to alter CL_sat_ abundance in cells.

**Figure 2:**
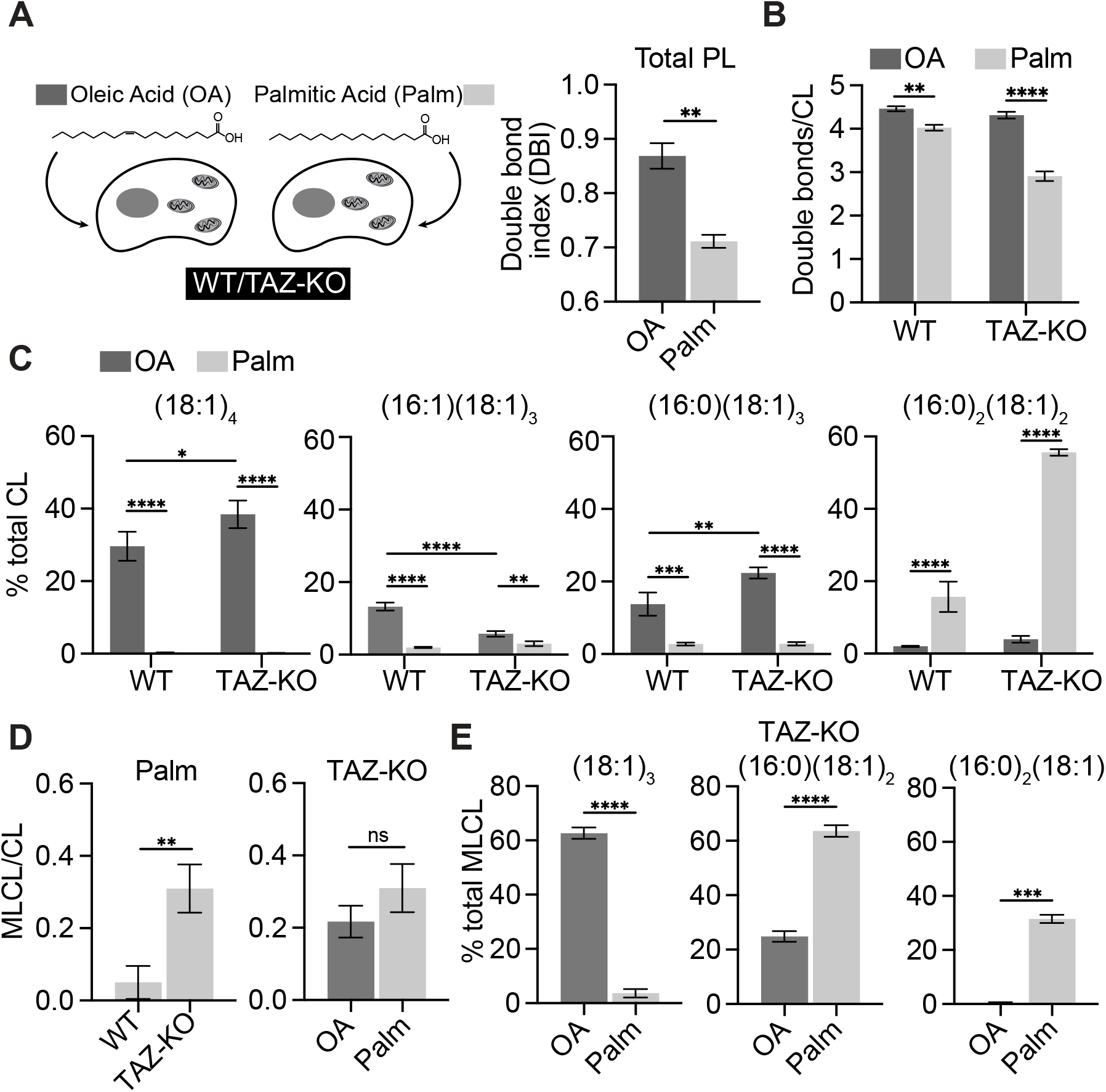
Manipulation of saturated CL abundance in C2C12 cells by fatty acid supplementation. (A) Exogenous addition of unsaturated and saturated fatty acids changes the double bond profile of the total PL pool in C2C12 myoblasts. Palm treatment decreased the double bond index (DBI) of the PL pool. DBI is defined as the mean number of double bonds in a given PL acyl chain. Cells were fed with either 100 μM of OA or Palm for 24 hours and subjected to lipidomics in biological triplicates (N=3). **P=0.0020 unpaired t-test of OA vs. Palm. Error bars indicate SD. (B) Treatment of TAZ-KO C2C12 myoblasts with 100 μM Palm results in a major increase in total CL saturation. Lipidomics analysis was performed on C2C12 cells in N=3 biological triplicates. **P=0.0010 WT OA vs. WT Palm. ****P<0.0001 for TAZ-KO OA vs. TAZ-KO Palm. ****P<0.0001 for POCL quantification for WT OA vs. WT Palm and for TAZ-KO OA vs. TAZ-KO Palm. P-values were obtained by two-way ANOVA analysis with Tukey’s correction for multiple comparisons. Error bars indicate SD. (C) Palm-treated TAZ-KO cells exhibit an increase in di-saturated CL species. ****P<0.0001, *P=0.0186 for WT OA vs. TAZ-KO OA for (C18:1)_4_ CL species, **P=0.0017 and 0.0013 for WT OA vs. TAZ-KO OA and TAZ-KO OA vs. Palm from C16:0(18:1)_3_ and (16:1)(18:1)_3_ respectively,***P=0.0003 WT OA vs. Palm for C16:0(18:1)_3_. P-values were obtained by two-way ANOVA analysis with Tukey’s correction for multiple comparisons. Error bars indicate SD. (D) Treatment of TAZ-KO cells with Palm does not significantly increase MLCL levels. An unpaired two-tailed t-test with Welch’s correction was used to determine statistical significance between OA and Palm TAZ-KO groups. Error bars indicate SD. (E) TAZ-KO cells treated with Palm exhibit increased incorporation of saturated fatty acids into MLCL. ****P<0.0001, **P=0.0070 for TAZ-KO OA vs. Palm for 56:6 MLCL species, ***P=0.0003 for TAZ-KO OA vs. Palm for (C16:0)_2_(18:1) MLCL species. P-values were obtained by an unpaired two-tailed t-test with Welch’s correction. Error bars indicate SD.

We next asked how changes in CL_sat_ would affect mitochondrial function. WT cells treated with OA or Palm exhibit indistinguishable, interconnected mitochondrial networks, as previously described (48). TAZ-KO cells treated with OA displayed interconnected mitochondrial structures, although with increased cell-to-cell heterogeneity compared to treatments in WT cells (Figure 3A). On the other hand, TAZ-KO cells treated with Palm displayed fragmented mitochondrial networks, a classical indicator of mitochondrial dysfunction (Figure 3A). Using an OXPHOS antibody cocktail, we also probed the abundance of key ETC and ATP synthase subunits (Figure 3B). Both OA and Palm-treated TAZ-KO cells had reduced abundance of ETC and ATP synthase markers compared to WT cells (Figure 3B), consistent with previous observations in untreated TAZ-KO cells (39, 40). However, Palm-treated TAZ-KO cells exhibited a complete loss of ATP-linked respiration compared to cells treated with OA, while WT cells did not exhibit altered respiration under either treatment (Figure 3C). Notably, while WT cells fed with Palm showed increased extracellular acidification rates (ECAR), an indicator of increased glycolysis (49–51), compared to OA cells, Palm-treated TAZ-KO cells exhibited both lower ECAR and oxygen consumption rates (OCR) compared to OA-treated cells, indicative of a quiescent metabolic state. Ultrastructural analysis of TAZ-KO cells treated with Palm displayed low cristae densities and an abundance of ‘onion’ or ‘empty’ cristae structures that resembled IMM structures in BTHS patients’ cells and animal models of BTHS (Figure 3D) (26, 41). Notably, these structures were absent from TAZ-KO cells treated with OA and treatments in WT cells (Figure 3D), suggesting that loss of TAZ sensitizes mitochondrial structure to Palm treatment.

**Figure 3:**
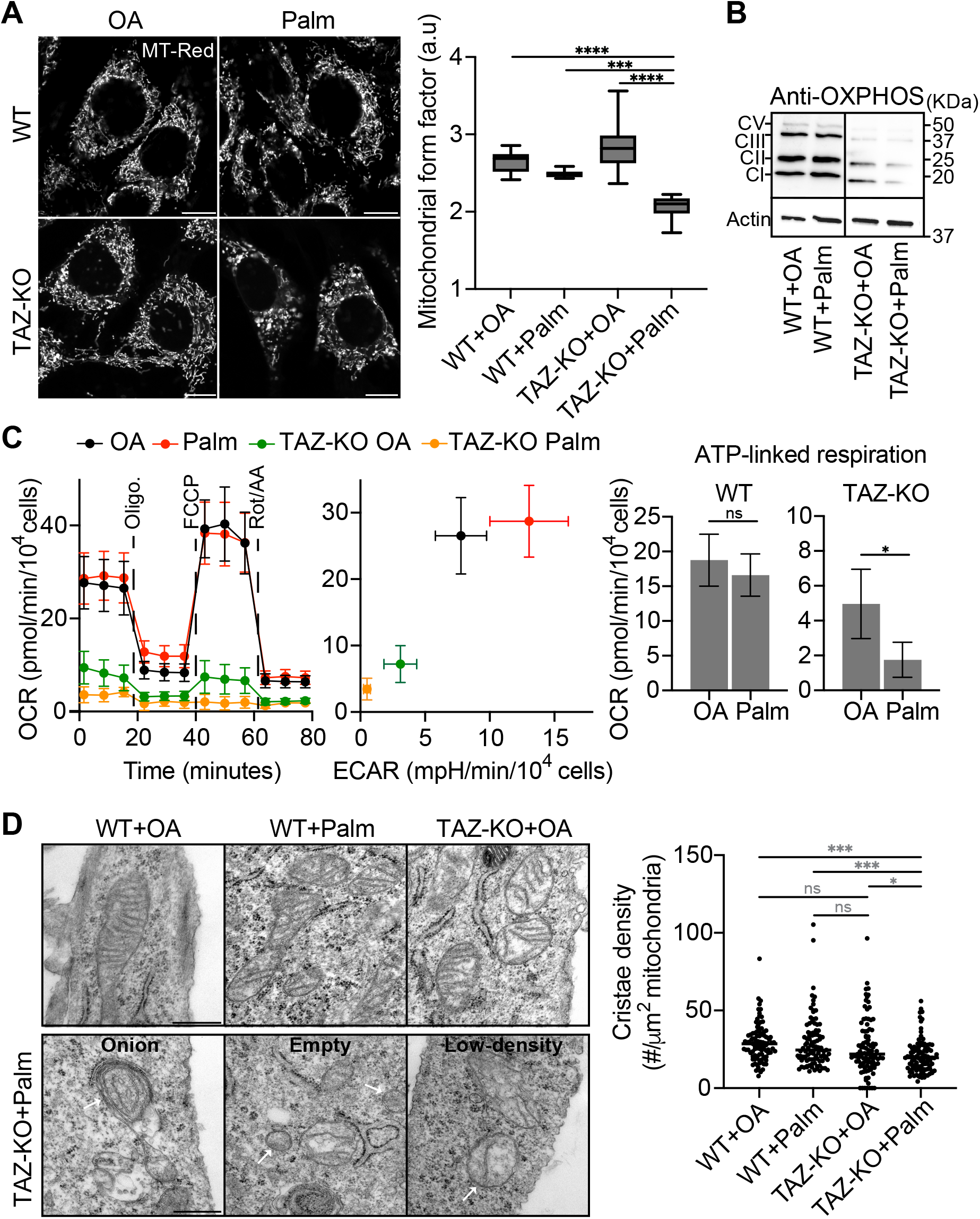
Accumulation of saturated CL species drives mitochondrial dysfunction. (A) TAZ-KO cells treated with Palm exhibit fragmented mitochondrial morphologies. Mitochondrial morphology was visualized by staining cells with 100nM Mitotracker red CMX-ROS. Scale bars, 10 μm. The mitochondrial form factor was quantified using the Mitochondria Analyzer plugin (81) from n=10 cells in N=3 biological replicates. Form factor values closer to 1 indicate spherical, fragmented mitochondrial networks while larger values indicate more complex topologies. ***P=0.0008 for WT Palm vs. TAZ-KO Palm, and ****P<0.0001 for WT OA and TAZ-KO OA vs. TAZ-KO Palm. P-values were obtained by two-way ANOVA analysis with Tukey’s correction for multiple comparisons. Error bars indicate SD. (B) Palm treatment in TAZ-KO cells results in a mild alteration to the abundance of OXPHOS complexes. 30 μg of whole cell extracts were subjected to SDS-PAGE and subsequently analyzed by western blot against an OXPHOS antibody cocktail. Actin served as a loading control. (C) Palm treatment of TAZ-KO C2C12 cells exacerbates defects in mitochondrial respiration. Seahorse respirometry of WT and TAZ-KO C2C12 cells treated with 100 μM of either OA or Palm. Wells containing 15,000 cells were treated in biological replicates (N>3) and subjected to a Seahorse XF cell mito stress test, involving a sequential addition of oligomycin, FCCP and antimycin A/rotenone. (B) Palm-treated TAZ-KO cells exhibit major defects in ATP-linked respiration. ATP-linked respiration is calculated by subtracting the oxygen consumption rate (OCR) after oligomycin-treatment from the basal respiration rate. (D) TAZ-KO cells treated with Palm lose cristae structure. Example electron micrographs of C2C12 myoblasts exhibiting cristae morphology after the indicated treatments. Scale bars, 250nm. Cristae density was quantified as the number of cristae tubules present per OMM area from (n>100) mitochondria. ***P=0.0002 WT OA vs. TAZ-KO Palm, ***P=0.0003 WT Palm vs. TAZ-KO Palm, *P=0.0408 TAZ-KO OA vs. TAZ-KO Palm. P-values were obtained by two-way ANOVA analysis with Tukey’s correction for multiple comparisons. Error bars indicate SD.

We hypothesized that the loss of mitochondrial structure in TAZ-KO cells upon treatment with Palm may be a result of lipidome-driven alterations to membrane physical properties, such as fluidity, which are sensitive to membrane acyl chain composition (3, 52, 53). The IMM in particular is enriched with unsaturated PLs, which contribute to its high membrane fluidity that is tightly coupled with respiratory state (53–55). We thus tested membrane fluidity in cells by staining with the IMM-targeted solvatochromic dye, Mito-Laurdan, and measuring Laurdan generalized polarization (GP) values by confocal microscopy (48). Lower GP values are characteristic of more fluid membranes whilst more rigid membranes exhibit higher GP values. WT cells treated with either OA or Palm exhibited similar mitochondrial GP values (Figure 4A-B), in line with our previous observations that IMM fluidity is resistant to mild (100 μM) Palm treatments, in contrast to other organelles like the ER (48). While TAZ-KO cells treated with OA exhibited similar GP distributions to WT cells, TAZ-KO cells treated with Palm showed increased GP values, indicating an increase in membrane ordering (Figure 4A-B). Thus, loss of Tafazzin sensitizes IMM fluidity to saturated fatty acid treatment.

**Figure 4:**
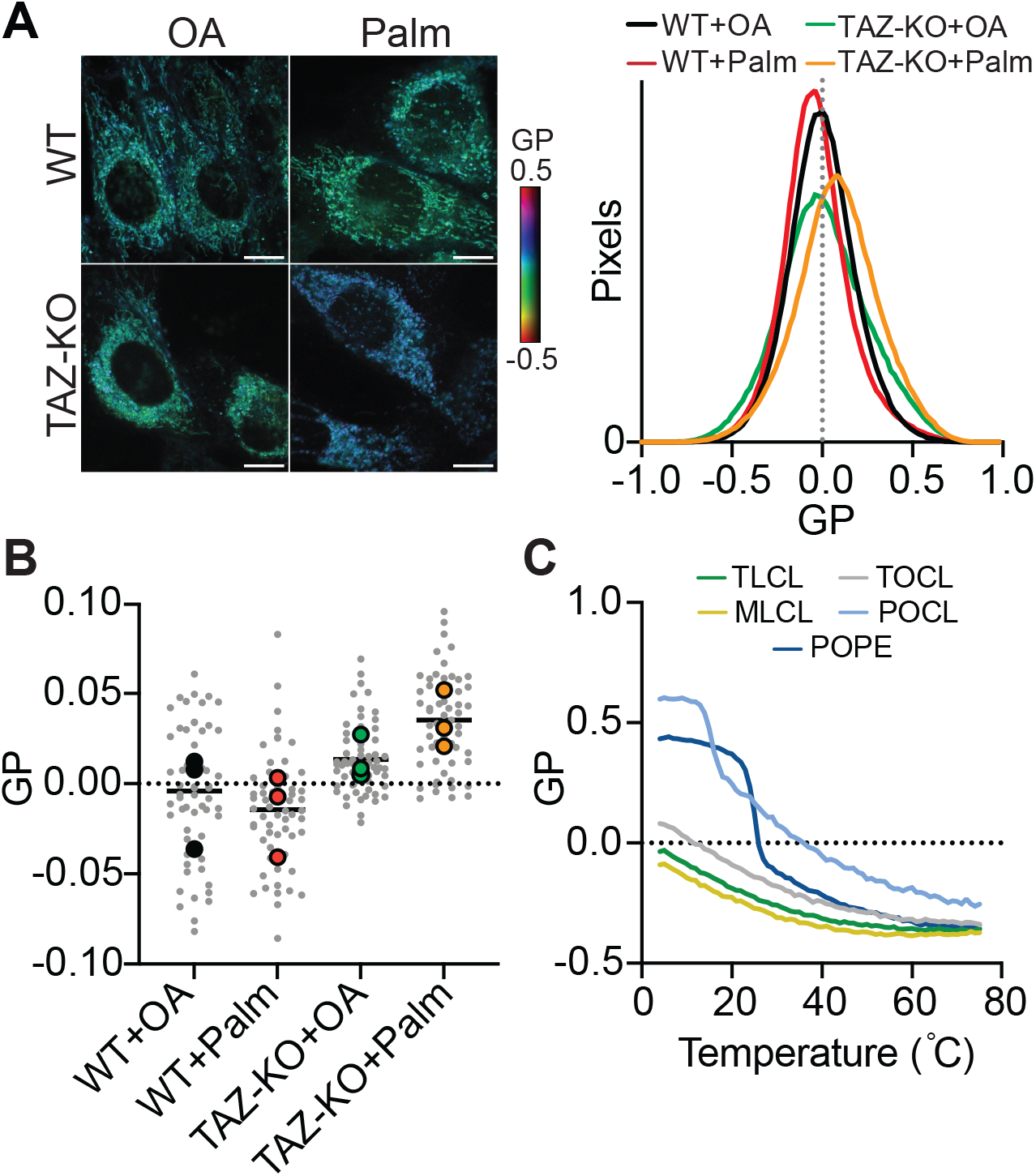
Saturated CLs increase membrane ordering of the IMM. (A) Mito-Laurdan (2.5 μM) fluorescence microscopy of WT and TAZ-KO C2C12 cells treated with 100 μM of either OA or Palm. Heatmap images are shown of Mito-Laurdan generalized polarization (GP) calculated from ordered and disordered channels. More positive GP indicates an increase in membrane order. Scale bars, 10 μm. Histograms of GP distribution by number of pixels of WT and TAZ-KO C2C12 cells treated with 100 μM of either OA or Palm. More positive GP indicates an increase in membrane order. (B) Mito-Laurdan mean GP of individual cells (light grey) and replicate means (colors) are shown of WT and TAZ-KO C2C12 cells treated with 100 μM of either OA or Palm. Values obtained from n=20 cells of N=3 replicates. P-values obtained by one-way ANOVA with multiple comparison correction (Dunnett’s test). (C) (A) POCL liposomes exhibit lower membrane fluidity compared to TOCL and MLCL. General polarization (GP) was determined after staining of liposomes with 2.5 μM C-Laurdan and analysis using a Cary fluorimeter. GP was calculated from emission intensities at 440 and 490 nm wavelengths after temperature ramps between 4-75°C. Tetralinoleic CL (C18:2)_4_, MLCL (C18:2)_3_, 1-palmitoyl 2-oleoyl phosphatidylethanolamine (POPE) was used as a control. High GP values indicate low membrane ordering. Above its melting temperature, TOCL liposomes show the highest GP values.

The reduced mitochondrial membrane fluidity in Palm-treated TAZ-KO cells supports the hypothesis that the loss of mitochondrial structure in these cells may be arising from alterations in membrane physical properties induced by changes in the lipidome. Previous MD and X-ray scattering studies have demonstrated reduced curvature sorting capabilities and increased lamellar-phase preferences for MLCL compared to tetra-oleoyl (C18:1)_4_ CL (TOCL) (34, 35, 56), however the physiochemical differences between CL_sat_ species like POCL and the well-studied TOCL have yet to be characterized. To address this, we first analyzed membrane fluidity properties of CL bilayers of altered acyl chain composition by staining liposomes with C-Laurdan (Figure 4C). Liposomes composed of unsaturated CL species TOCL, TLCL and (C18:2)_3_ MLCL, exhibited high membrane fluidity (Figure 4C). In comparison, POCL formed more ordered bilayers, with comparable GP profiles to those of (16:0 18:1) phosphatidylethanolamine (POPE), a high-melting temperature (T_m_) phospholipid (Figure 4C). The T_m_ of lipids are often indicative of their relative fluidity, with higher fluidity membranes exhibiting lower T_m_ (57, 58). While TOCL and DOPC did not melt above 0°C, POCL exhibited a melting temperature of 14.2°C (Figure S2), further implying increased membrane ordering.

In addition to membrane fluidity, alterations to acyl chain composition impact lipid spontaneous curvature (59), which can contribute to membrane curvature (3, 60) that is especially relevant in the highly-folded IMM. CL shows a cation-dependent curvature that could be relevant to its organization in the IMM (4, 6). We thus asked the role of CL saturation on this property. Lipid spontaneous curvature, *c*_0_, is traditionally measured by small angle X-ray scattering (SAXS), where lipid suspensions yield radially symmetric scattering patterns that correlate to profiles with distinct peaks that are used to identify lipid phase and its dimensions (61). Values for *c*_0_ are calculated from a nonbilayer inverted hexagonal phase, H_II_, where *c*_0_ is the inverse of the H_II_ tubule radius approximated at the neutral plane (3). Most lipids, such as TOCL, do not readily form H_II_ phases, and are instead included as ‘guests’ in a mixture with an H_II_-forming ‘host’ lipid such as dioleoyl-phosphatidylethanolamine (DOPE) (Figure 5A). While previous studies have determined *c*_0_ values for TOCL (62), *c*_0_ of other CL species, such as POCL have yet to be determined. We thus utilized SAXS to experimentally determine the *c*_0_ of POCL in comparison to TOCL (Figure 5A). We measured POCL to have a *c*_0_ of −0.008Å^−1^, comparable to the low-curvature POPC (−0.0029Å^−1^) and reduced in magnitude compared to TOCL (−0.0176Å^−1^). Our values for TOCL were in agreement with those previously measured under similar ionic conditions (5mM Ca^2+^) (62). Thus, POCL is a low-curvature CL species.

**Figure 5:**
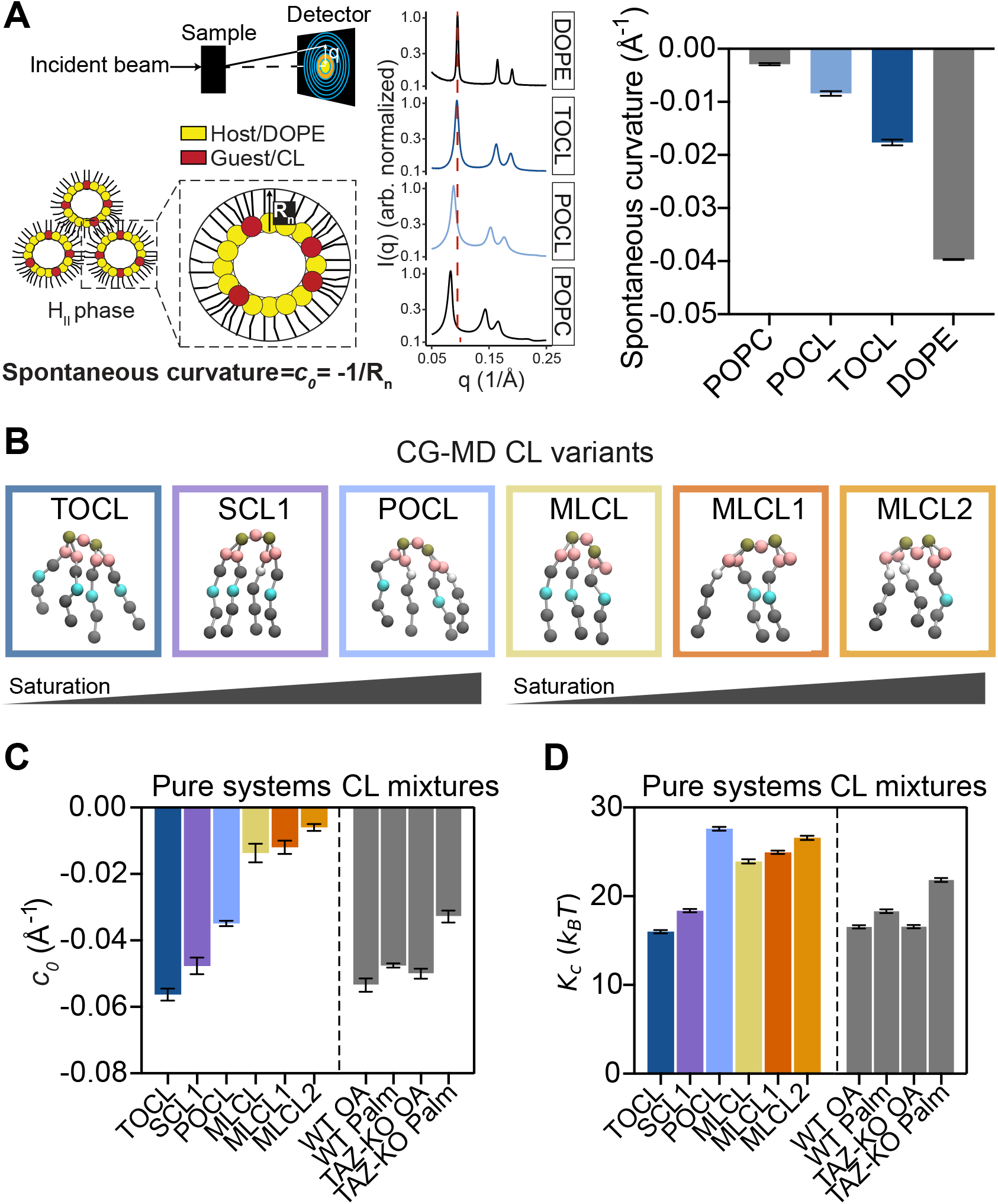
Altered mechanical properties of saturated CLs. (A) POCL is a low curvature CL species. Spontaneous curvatures values at 35°C were determined using small angle X-ray scattering (SAXS) on hydrated lipid suspensions that are diagramed to the left. Representative SAXS profiles of the H_II_ phase of 100% DOPE, and hosted TOCL (10%), POCL (10%) and POPC (10%) in DOPE (90%) are shown. The dashed line indicates the scattering angle of the first H_II_ Bragg peak in pure DOPE. Shifts to the left indicate widening of H_II_ tubules, corresponding to increases in *c*_0_. All samples were relaxed in 12% (w/w) tricosene. Extrapolated curvatures (*c*_0_) are plotted to the right. (B) Martini3 models for CL_sat_ species in comparison to previously employed TOCL and MLCL components. (C) Extrapolated curvatures (*c*_0_) from CL bilayers. The pure systems reflect unicomponent membranes of CL and MLCL species with increasing saturation shown in B. The mixtures reflect combinations of CL species based on lipidomic profiles of C2C12 WT and TAZ-KO cells supplemented with either Palm or OA. (D) Corresponding bending moduli (*K*_c_) for the systems shown in C. Like for *c*_0_, *K*_c_ is most affected by saturated CL species like POCL and the CL composition found in TAZ-KO Palm cells.

Molecular dynamics (MD) is a powerful tool for investigating how lipid structural changes translate membrane mechanical properties. Previous MD studies have demonstrated the reduced propensity for MLCL to localize to regions of high curvature compared to TOCL (34), however analogous analyses on CL_sat_ have been lacking. To answer this, we built and validated an array of CL molecules with differing acyl chain compositions using coarse-grained (CG) MD using the Martini 3 forcefield. We chose CG-MD because height fluctuation analysis of large, undulating bilayers can be used to derive membrane bending rigidity (*K*_c_), a key property for formation of high-curvature membrane structures like in cristae. CG-MD simulations can also be used to conduct large-scale simulations of realistic IMM architectures (63, 64). To investigate properties of CL_sat_, we developed two new models for CL molecules with saturations on each acyl chain, SCL1 and POCL, and two new models for MLCL with saturations on each acyl chain, MLCL1 and 2 (Figure 5B, Figure S3). These models were then used in large (40×40 nm) and small-scale (15×15 nm) bilayers that were run in parallel with identical compositions. The former was used to derive *K*_c_, while the latter was used to calculate lateral pressure profiles (LPP). The first moment of the LPP is equal to *-K*_c_*c*_0_, where *c*_0_ is the spontaneous curvature (65). By measuring *K*_c_ independently in the large systems, values for *c*_0_ in the smaller systems of the same composition can be estimated.

Our simulations indicated that CL with two saturated acyl chains, the CL_sat_ species POCL, can especially affect biophysical properties relevant for membrane bending. Values derived for spontaneous curvature and bending rigidity are plotted in Figures 5C and 5D. Overall, CG-MD of CL-containing membranes showed low *K*_c_ values, as observed by previous studies (6, 66), which led to larger, more negative *c*_0_ values than measured by experiments. Nonetheless, we observed a consistent trend in the role of saturation on these parameters. While SCL1 showed modest changes in *c*_0_ (Figure 5C) and *K*_c_ (Figure 5D) compared to TOCL, these parameters were markedly altered in POCL. Increasing saturation of MLCL also changed both these properties, albeit more gradually, with 2 saturated acyl chains on MLCL2 resulting in a *c*_0_ close to 0. Although we did not measure *K*_c_ of POCL experimentally, its reduced fluidity and T_m_ compared to TOCL (Figure 4C) is consistent with a decrease in *K*_c_ due to this parameter’s proposed quadratic scaling to increases in membrane thickness (67). Notably, POCL exhibited a larger *K*_c_ than all tested MLCL systems, exemplifying that POCL specifically contributes to membrane stiffness (Figure 5D).

CG-MD provides an opportunity to sample complex lipid mixtures that are found in cells. We thus simulated systems containing compositions of multiple CL and MLCL species extracted from our lipidomics data (Table S1). From these simulations, we extracted *K*_c_ and *c*_0_ parameters in a pipeline identical to the pure CL systems. We observed only minor changes in *K*_c_ and *c*_0_ parameters between WT OA, WT Palm and TAZ-KO OA mixtures (Figure 5C-D). In contrast, TAZ-KO Palm, which accumulates the most POCL, showed the most reduced *c*_0_ and the largest increase in *K*_c_. Thus, our *in vitro* and *in silico* results indicate an altered physiochemistry of POCL molecules compared to their unsaturated counterparts. They predict that experimental combinations of Palm treatment and Tafazzin loss causes the CL pool to become one with reduced intrinsic curvature and increased stiffness. These parameters could in turn be relevant for the formation of high-curvature cristae that are lost in Palm-treated TAZ-KO cells.

## Discussion

In this study we identify that CL_sat_ species alter membrane physical properties and can drive mitochondrial dysfunction when they accumulate in a C2C12 skeletal myoblast model of BTHS. CL_sat_ and MLCL species both accumulate in BTHS patients’ (47); while MLCL has been shown to lack the curvature-producing capabilities of unsaturated CL (32, 34) and reduce IMM protein stability (31, 33), analogous mechanisms by which CL_sat_ molecules impact mitochondrial structure and function have been lacking. Separating contributions of MLCL and CL_sat_ as drivers of mitochondrial dysfunction has been challenging. Low levels (25 μM) of Palm do not significantly alter CL acyl chain profiles while high levels (>200 μM) increase CL_sat_ but have pleiotropic effects (68), and are known to induce ER stress (69). TAZ-KO cells lost cristae (Figure 3) when fed with 100 μM Palm, a treatment which does significantly affect organellar morphology and function in WT cells (48).

The conserved remodeling pathway serves to enhance the unsaturation of CL compared to other lipid classes, like PG and CDP-DAG that are its direct metabolic precursors. Here, TAZ-KO+Palm cells accumulated di-saturated CL’s (16:0 18:1)_2_ (POCL) as the major CL species (Figure 2), with other changes in CL species matching WT cells treated with Palm. Our results extend our previous observations in yeast that the functionality of CL remodeling is dependent on the surrounding lipid environment (44), as WT cells treated with Palm and TAZ-KO cells treated with OA still retained a high level (DB∼4) of unsaturation in CL. The effect of CL_sat_ accumulation on mitochondrial function and cristae structure raises several questions regarding potential roles of CL_sat_ as a signal of mitochondrial damage. Previous studies have shown that exogenous addition of CL_sat_ species induces inflammation, while unsaturated CL species inhibit inflammatory responses (70, 71). The externalization of CL is known to induce macrophage-mediated mitochondrial clearance (72), however it remains to be studied how the presentation of CL is altered in BTHS, where patients exhibit neutropenia (73), potentially from excessive clearance of neutrophils by macrophages.

To the best of our knowledge, this study presents the first analysis of the membrane physical properties of physiologically relevant CL_sat_ species. Using TAZ-KO cells fed with Palm, we characterize the biophysical properties of POCL (16:0 18:1)_2_, the major CL species that accumulates in BTHS patient lymphoblasts alongside MLCL (47). In experimental analyses, we find that POCL exhibits a reduced fluidity and magnitude of intrinsic curvature *c*_0_ compared to TOCL (Figure 4-5). This was complemented by *in silico* analysis of a wider range of compositions. Our results suggest that CL_sat_ species with two saturated chains, like POCL, have particularly affected properties. These include a reduced magnitude of *c*_0_ and membranes with an increased bending moduli *K*_c_. Both *c*_0_ and *K*_c_ are central parameters in the Helfrich model of membrane bending (74), and these increases would increase the free energy to form high-curvature structures in the IMM. A systematic analysis of MLCL has also been shown previously to have positive *c*_0_ by atomistic MD, and MLCL-containing bilayers buckle at higher lateral pressures compared to TOCL by CG-MD, indicating limited propensity for shape deformations (7, 34). These results imply that both CL_sat_ and MLCL could act to alter mitochondrial physiochemical properties and thereby contribute to defects in IMM structure and function. Notably, a loss of high-curvature cristae ultrastructure was originally observed in the bone marrow of BTHS patients (26), an effect that we can recapitulate in C2C12 cells only upon accumulation of both MLCL and CL_sat_ under Palm-fed conditions. The sensitivity of CL’s properties to acyl chain saturation also provides a rationale for why a CL-specific remodeling pathway evolved in eukaryotes (75).

An open question is how CL_sat_ accumulation could also directly affect the stability of ETC complexes and cristae-shaping proteins themselves. A recent study demonstrated equivalent binding capabilities of optic atrophy protein 1 (OPA1) to POCL and 25% MLCL membranes, both of which had reduced binding capabilities compared to tetralinoleic CL (18:2)_4_ but surprisingly equivalent binding to TOCL (36). Fully saturated CL species such as TPCL (16:0)_4_, exhibit low binding capabilities to *cytochrome c* and OPA1 (36, 76). However, it is important to note that TPCL is not detected in BTHS patients’ lipidomics or in Palm-fed TAZ-KO cells, and exhibits gel-like phase properties at 37°C and is therefore unlikely to be physiologically relevant (77, 78). Thus, the effect of POCL on ETC stability is physiologically relevant and warrants further study. Altered functions and/or interactions of cristae-shaping proteins in response to CL_sat_ species could represent an additional mechanism by which inner membrane structure is affected by loss of the CL remodeling pathway.

Several therapeutic strategies to mitigate mitochondrial dysfunction in BTHS have focused on ameliorating the MLCL:CL ratio, which has been achieved through iPLA_2_ inhibition by BEL or by inhibition of ABHD18 (17, 19, 20). However, these treatments do so by inhibiting the initiation of CL remodeling. Thus, they would cause accumulation of CL_sat_ species (18, 20), which may still impair mitochondrial function, preventing a complete reversal of the BTHS phenotype. Our previous results also suggest that mitochondrial integrity in BEL-treated cells are particularly susceptible to Palm supplementation (3), indicating that further studies should test the effect of the surrounding lipid environment in cells treated with ABHD18 inhibitors, such as ABD646 (20). When CL remodeling is lost, the acyl chain composition of the remaining de novo-synthesized CL is more sensitive to precursor phospholipids and local changes to fatty acids dictated by diet and metabolism (Figure S1). This dynamic is not necessarily captured in cultured cells grown in serum enriched in exogenous unsaturated fatty acids and could be relevant for tissue- and patient-specific phenotypes that characterize remodeling disorders like BTHS. For BTHS, approaches that combine inhibition of MLCL formation and alterations of fatty acid metabolism to reduce CL_sat_ may provide an improved therapeutic strategy.

## Materials and Methods

### Mammalian cell culture

C2C12 skeletal myoblast cells were maintained in DMEM (Gibco) supplemented with 10% fetal bovine serum (FBS, Gibco) and 1% antibiotic/antimycotic and grown in 37°C incubators containing 5% CO_2_. For fatty acid treatments, cells were supplemented for 24 hours with either 100 μM palmitate (Palm) or oleate (OA), prior to analysis. Palm was prepared as previously described (80): Sodium palmitate or sodium oleate (Thermo Fisher) was complexed to fatty acid free Bovine Serum Albumin (BSA). Fatty acids were dissolved 1:1 in a mixture of 0.1 mM NaOH in Hanks’ balanced salt solution (HBSS) and ethanol to a concentration of 200 mM, heated to 65°C, and then added to a 4.4% BSA HBSS solution that had been heated to 37°C such that the final fatty acid concentration was 2 mM in a 3:1 molar ratio with BSA. The mixture was then stirred vigorously and incubated at 37°C for 1 hour. The Palm:BSA mixture or 4.4% BSA vehicular control was diluted into complete DMEM containing 10% FBS to a concentration of 100 μM and filtered through a 2 μm polyethersulfone (PES) filter (Fisher Scientific) before use. Cells were seeded and then grown with complete DMEM containing Palm complexed to BSA and subjected to analysis after 24 hours. The treatment time was selected as there are no observable defects to organelle morphology, but an increase in esterified saturated fatty acids (80). Cells were routinely tested for mycoplasma and not detected.

### Confocal microscopy

Live cell microscopy was conducted using Plan-Apochromat 63x/1.4 Oil DIC M27 objective on the Zeiss LSM 880 with an Airyscan detector; image acquisition and processing was performed with ZEN software using default processing settings. To assess mitochondrial morphology, 15,000 cells were seeded into 8-well coverglass-bottom chambers (Nunc Lab-Tek) pre-treated with 5μg/mL human plasma fibronectin (Thermo). Cells were treated with fatty acids and grown for 24 hours prior to staining with 100nM Mitotracker red CMX-ROS for 20 minutes, three washes with HBSS, and resuspension in clear DMEM without phenol red before imaging using a 561nm laser line at 0.2% power on the Zeiss LSM 880 with Airyscan. Mitochondrial morphology was assessed from n>10 cells in biological triplicates (N=3) using the Mitochondria-Analyzer plugin on ImageJ (81).

Measurements of mitochondrial membrane fluidity were performed essentially as previously described (48). WT and TAZ-KO cells were seeded in 8-well chamber slides (LabTech) and treated with either 100 μM of Palm or OA for 24 hours. Cells were washed once with HBSS prior to staining with 2.5 μM of Mito-Laurdan for 20 minutes in DMEM clear media with glucose and without phenol red. Cells were then washed once with HBSS and subsequently washed with DMEM clear media containing 10% FBS for 6 minutes per wash. Cells were then imaged in DMEM clear media containing 10% FBS. Mito-Laurdan was excited using an 405nm laser line and emission was collected using an airyscan detector with a 420-480nm bandpass filter. Images were processed using ImageJ and GP analysis was performed using a custom-built script on n=20 cropped cells in triplicates (N=3).

### Lipidomics

C2C12 myoblast cells were seeded and grown in 6-well plates with the indicated treatments for 24 hours before being washed with HBSS and scraped. Cells were initially homogenized into 1ml of HBSS. A mix of deuterated internal standards for each lipid class (Equisplash, Avanti) was added to 250ul of the homogenate. All lipids were extracted by modified BUME and measured according to the method described previously (82) The lipids were measured using a Q-Exactive orbitrap mass spectrometer (Thermo Fisher Scientific) interfaced with a Vanquish UHPLC (Thermo Fisher Scientific). A Waters T3 1.6uM 2.1mm x 150mm column was used for chromatographic separation using a step gradient from 25% buffer A (10mM ammonium formate and 0.1% formic acid in water) to 100% buffer B (70/30 isopropanol/acetonitrile with 10mM ammonium formate and 0.1% formic acid) over 35 min. Flow rate was set at 0.3 mL/min. Lipid analytes were analyzed using a data dependent acquisition (DDA) Top N scan of 8 with an NCE of 30 in negative mode. All ions in the mass range of 200-2000 m/z were monitored. MS1 resolution was set at 70,000 (FWHM at m/z 200) with an automatic gain control (AGC) target of 1e6 and Maximum IT of 200 ms. MS2 resolution was set to 17,500 with an AGC target of 5e4, fixed first mass of 80 m/z, and Maximum IT of 50 ms. The isolation width was set at 1.2; Dynamic Exclusion was set at 3s. Lipid Identification and Quantification was done with Lipidomic Data Analyzer Software (83).

Lipidomic Data Analyzer Software identified CL molecular species in negative mode. The precursor fragment used was [M-H]^−^. Pure TOCL 18:1/18:1/18:1/18:1, POCL 16:0/18:1/16:0/18:1 and MLCL 18:2/18:2/18:2 (Avanti Polar Lipids) was used as internal standards. The PG headgroup is identified by the fragment [C_3_H_6_O_5_P]^−^ at m/z 152.99 m/z. The fragments used for the sn2 chain identification are the four carboxy fragments. For example, CL 16:0/18:2/18:2/18:2 gives a fragmentation pattern of: precursor [C_75_H_141_O_17_P_2_]^−^ at 1376.97 m/z, PG headgroup fragment [C_3_H_6_O_5_P]^−^ at m/z 152.99 m/z, and carboxy fragments [C_16_H_31_O_2_]^−^ at 255.23 m/z and [C_18_H_31_O_2_]^−^ at 279.23 m/z. Noise filtering and fragmentation intensity rules were established to prevent false positive identification.

### Respirometry

For analysis of respiration in C2C12 cells, cells were treated for 24 hours prior to analysis on the Agilent Seahorse XF pro (Agilent). 15000 cells were seeded in biological replicates (n=4) into 96 well Seahorse cell culture microplates (Agilent) pre-treated with fibronectin (5 μg/mL). Samples were analyzed using the Seahorse Cell Mito Stress Test, with sequential addition of Oligomycin (1.5 μM), FCCP (1 μM) and a Rotenone/Antimycin A mixture (0.5 μM). Analysis was performed on respiration rates after treatment with inhibitors to readout basal, ATP-linked, maximal and proton leak respiration rates. After the assay, cells were trypsinized and mixed with trypan blue and cell-counted to normalize respiration rates.

### Electron microscopy

C2C12 cells were first cultured to confluency in each treatment condition prior to fixation in a pre-warmed 4% glutaraldehyde solution for 30 minutes at room temperature. Samples were then re-fixed and embedded prior to imaging using a JEOL 1400 transmission electron microscope. ImageJ software was utilized to measure cristae tubule lengths and OMM areas from thin section images. For quantification of cristae density, the number of cristae within n=100 mitochondria were counted in each condition and normalized to the OMM area.

### Liposome preparations

Pure phospholipids were purchased from Avanti Polar Lipids as chloroform solutions of known concentrations. Constituents of lipid mixtures were measured using Wiretrol II glass pipette tubes (Drummond) attached to micropipettors. For analysis of membrane fluidity and melting temperature, lipid dispersions were made by drying lipid films under N_2_ and high vacuum for 30 min, then resuspending them in buffer (20 mM HEPES, 2 mM EDTA, pH 7.4) to concentrations of approximately 1 mM lipid. Samples were then subjected to 4 freeze-thaw cycles and subsequently extruded using a hand-held device (Avanti Polar Lipids) incorporating a 200 nm polycarbonate track-etch filter (Whatman) at room temperature. For analysis of membrane fluidity, the resultant unilamellar vesicles were stained with 2.5 μM C-Laurdan and tumbled for 30 min at room temperature prior to analysis on a Cary Eclipse Fluorimeter (Agilent). Generalized polarization (GP) of C-Laurdan was calculated from emission intensities at 440 and 490 nm after excitation at 340nm with 5nm excitation and emission slits. Emission intensities were acquired from a temperature ramp between 4-75°C at a rate of 1°C/min. For determination of lipid melting temperatures, differential scanning calorimetry (DSC) was performed on 1mM resuspended lipid dispersions. Prior to analysis, both lipid dispersions and buffer were degassed for 15 min prior to loading on DSC 2500 (TA instruments). Samples and buffers were subjected to heating scans between 0-100°C. TA Nanoanalyze software was used to visualize thermograms necessary for determination of lipid melting point.

### Small angle X-ray scattering sample preparation and analysis

Lipid samples were prepared essentially as previously described (84), 2.5 mg of pure lipids, including 12% (w/w) (Z)-9-Tricosene, were dissolved in 50 μL of cyclohexane (Fisher Chemical) and transferred to 1.5 mm ID capillary melting point tubes (Kimble Chase). Tubes were then frozen at −80°C and lyophilized for 1 hour prior to rehydration by addition of 10 μL of vacuum-degassed miliQ water. Rehydrated samples were subjected to five freeze thaw cycles in an isopropanol-dry ice bath and homogenization with a 20 μL Wiretrol II microdispenser (Drummond Scientific). The bottom of the tube was scored and cut before transferring the suspension to a disposable sample holder. The suspension was settled to the bottom of the sample holder by gentle centrifugation at 500 RCF for 5 min at 4°C). Full hydration was ensured by addition of 5 μL of degassed water to the top of the sample. High vacuum grease (Dow Corning) was added to the top of the water to seal the sample prior to analysis.

SAXS data was collected essentially as previously described (84, 85) on beamline 7A at the Center for High-Energy X-ray Sciences using a custom-built temperature-controlled pressure cell (86). The experimental parameters were as follows: photon energy was 14 keV, spot size was 200 x 250 µm (W x H), and scattering data were collected for 0.005 Å-1 ≤ q ≤ 0.7 Å-1 using an EIGER 4M detector (Dectris) within the beamline vacuum. 1 s exposures were taken at a flux of 1.8-3.6×10^11^ photons/s. Three initial exposures were used to test each sample for radiation sensitivity, then pressure or temperature up- and down-sweeps were made starting at the lowest temperature or pressure and proceeding to the highest. The sample was equilibrated for at least 60s per 50 bar of pressure change. Because photon scattering by the lipid dispersions was strong and background was low (as assessed by shooting a sealed capillary containing the same degassed water used to hydrate the lipids), no background subtraction was used. SAXS images were integrated using BIOXTAS RAW software (87).

### Measurement of phospholipid monolayer curvature

For *c*_0_ measurement, pure lipids (Avanti Research) were prepared as hosted mixtures in DOPE at a molar ratio of 10 mol%:90 mol% guest:host for HPSAXS, with the addition of 12% w/w 9(Z)- tricosene, as described above. 10 mol% MLCL, POCL, and TOCL samples contained 5mM Ca^2+^ and 33 mM Na^+^ as a result of the anionic lipids being utilized as sodium salts. Neutral-plane *c*_0_ was calculated by using a constant distance correction of 3.94 Å, derived from global fitting of the DOPE H_II_ scattering profile (88). Guest lipid spontaneous curvature was calculated from a weighted average of the total guest in host system curvature.

### Molecular dynamics analyses

The membrane model for each mixture-system was constructed by creating a symmetric bilayer whose lipid compositions were derived from lipidomics. The corresponding homogeneous systems were also constructed for each lipid type (Figure S3): TOCL, SCL1, POCL, MLCL, MLCL1, MLCL2. If multiple isomers were possible, we chose the following: MLCL lacks chain 4 (SN1); MLCL1 lacks chain 4 and has chain 2 (SN1) fully saturated; MLCL2 lacks chain 4 and has chains 1 (SN2) and 2 (SN1) fully saturated. All unsaturated chains correspond to oleic acid (18:1) and all saturated chains correspond to palmitic acid (16:0) in agreement with the biosynthesis of these lipids. As the only lipid of this set in the Martini3 database is TOCL, we created the rest of the variants by changing beads and bonds type accordingly (Figure S3).

We employed a coarse-grained molecular dynamics (CG-MD) simulation framework using the Martini3 force field (89). For each composition, two system sizes were considered: one of approximately 40 × 40 × 25 nm and the other one of 15 × 15 × 25 nm. These configurations were assembled using the insane.py utility (90). All simulation boxes were hydrated and neutralized with Na^+^ following the standard insane.py workflow (Table S2). Energy minimization and equilibration steps followed procedures adapted from the CHARMM-GUI Martini Maker (91). From configurations generated by insane.py, systems were minimized using conventional steepest descent minimization for 2000 steps. Minimized systems then undergo restrained equilibration where harmonic restraints holding the lipid head groups in place are relaxed from 200–50 kJ mol^−1^ nm^−2^ with concomitant increase of timestep from 2–20 fs over a series of steps. All equilibration steps are performed in the NPT ensemble at 1 bar. The first equilibration step was performed at 340 K to prevent gel formation, as lower temperatures are known to promote more ordered phases within the Martini3 force field (89). From the second equilibration step and onwards, the temperature was set at 310 K to mimic the physiological relevant temperature. To verify the correct equilibration of all systems, we calculated the area per lipid across the equilibration steps and production, which converged by frame 2000 (Figure S4).

Production trajectories were run for 5 μs using a 20 fs timestep. The small systems were simulated for 3 μs, followed by an additional 300 ns using a 5 ps output interval necessary for the stress analysis. Reported simulation lengths represent the actual unscaled Martini simulation times rather than post-processed, time-corrected equivalents. During all equilibration steps and production, pressure control relied on a C-rescale semi-isotropic barostat (τ_p_ = 4 ps; compressibility 3 × 10^−4^ bar^−1^) set to 1 bar (92). Electrostatic interactions were computed via the reaction field method with a 1.1 nm cutoff and ϵ_r_ of 15 (93), and the temperature was held using the stochastic velocity-rescaling thermostat with τ_t_ set to 1.0 ps (94).

The mechanical properties of the membranes were estimated from the simulations following the same protocol as described in previous work (6). Briefly, the bending modulus was estimated from each large-box simulation by analysis of the height fluctuation spectra. The spontaneous curvature was estimated from the moment of the lateral pressure profile and substituting the estimated bending modulus for each small-box simulation. All codes for simulation set up and analysis can be found online at (95). Software versions used are documented in Table S3.

## Supporting information

Combined Supplementary Information

## Acknowledgements

Miriam Greenberg generously provided C2C12 WT and TAZ-KO cell lines. Jacob Winnikoff provided assistance with SAXS sample preparation, measurement, and analysis. Neal Devaraj and Shane Douty contributed equipment and expertise for DSC measurements. Beamline scientists Qingqiu Huang, Estella Yee, and Richard Gillilan provided assistance at the Center for High-Energy X-ray Sciences (CHEXS). Lipid analysis was performed at the LIPID MAPS Lipidomics Core at UC San Diego. Molecular dynamics simulations were run on hardware hosted by the Triton Shared Computing Cluster (79). We are also grateful to the UCSD Physics Computing Facility for computational resources.

## Funding

Research was supported by the National Institute of General Medical Sciences (R35-GM142960 to I.B., R35-GM139641 to E.A.D., T32--GM008326C to K.V., T32-GM133351 to C.S.), the National Heart, Lung, and Blood Institute (R01HL157115 to X.F), National Science Foundation (MCB-2046303 to I.B.), the Department of Energy (DE-SC0022954 to I.B.), the American Heart Association (25EIA1369578 to X.F), the Allen Family Philanthropies, and the Barth Syndrome Foundation. Data acquisition at CHEXS is supported by the NSF award DMR-2342336, and the MacCHESS resource is supported by NIGMS award 1-P30-GM124166.

## Author contributions

K.V. and I.B. conceived the study. K.V, D.M, E.K, A.M.W, and A.A carried out experiments and analysis. C.S., C.M.S., and C.T.L designed and analyzed simulations.

## Conflict of interest

The authors declare that they have no conflicts of interest.

## Notes

### Competing Interest Statement

The authors have declared no competing interest.

