## Supplementary material for "Saturated cardiolipins are potent disruptors of inner mitochondrial membrane structure and function": Combined Supplementary Information

This file contains:

Figures S1-S4
Table S1-S3

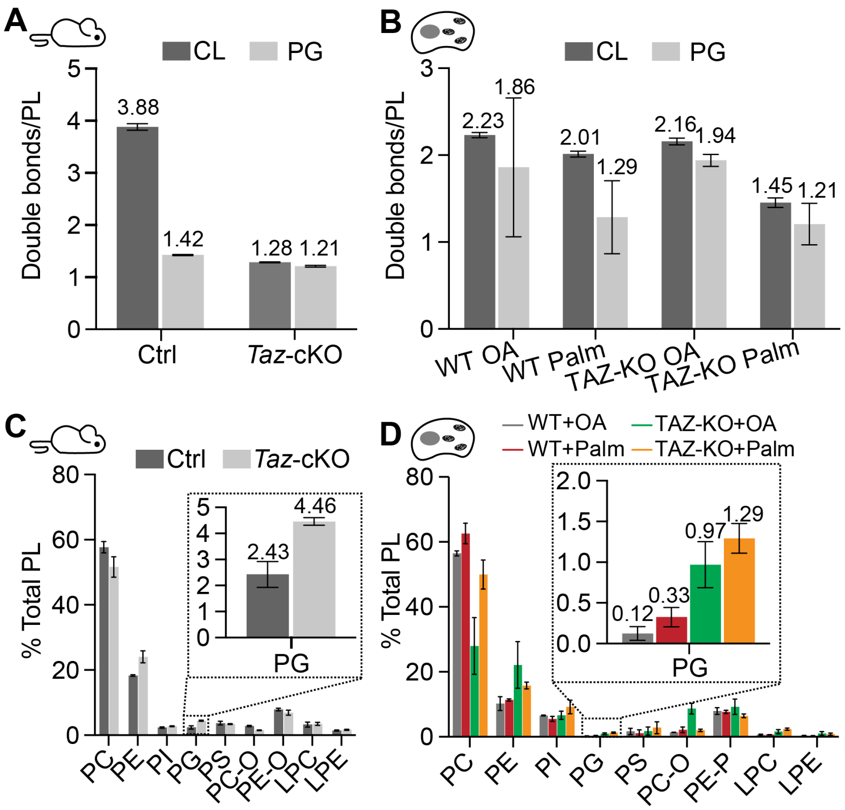


**Figure S1:** Lipidomics analysis of PG species from mouse hearts and C2C12 cells

(A) & (B) *Taz*-cKO mouse hearts and TAZ-KO C2C12 cells treated with Palm exhibit similar chain-adjusted CL and PG saturation levels. CL molecules contain 4 acyl chains while PG molecules contain 2 acyl chains, so double bonds/PL are shown for chain-adjusted CL molecules (double bonds/CL divided by 2). Lipidomics analysis of PG species from control and *Taz*-cKO mouse hearts, n=3-4 mice per group, and WT and TAZ-KO C2C12 cells upon treatment with 100 µM of Palm or OA in N=3 biological triplicates. Error bars indicate SD. (C) & (D) *Taz*-cKO mouse hearts and TAZ-KO C2C12 cells exhibit increased levels of PG. Total PL abundances for control and *Taz*-cKO mouse hearts, n=3-4 mice per group and WT and TAZ-KO C2C12 cells upon treatment with 100 µM of Palm or OA in N=3 biological triplicates. Error bars indicate SD.

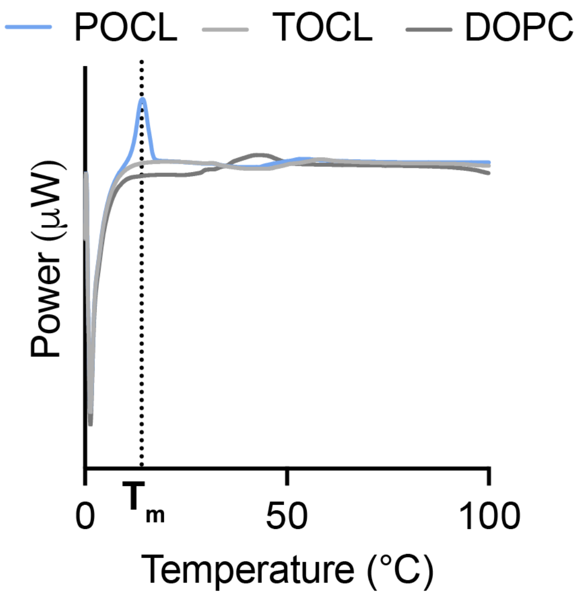


**Figure S2:** The melting temperature of POCL measured by Differential scattering calorimetry (DSC). POCL liposomes yield a 14.2 °C melting temperature. No melting was observed for TOCL or DOPC in the tested range. DSC was performed on 1 mM lipid dispersions.

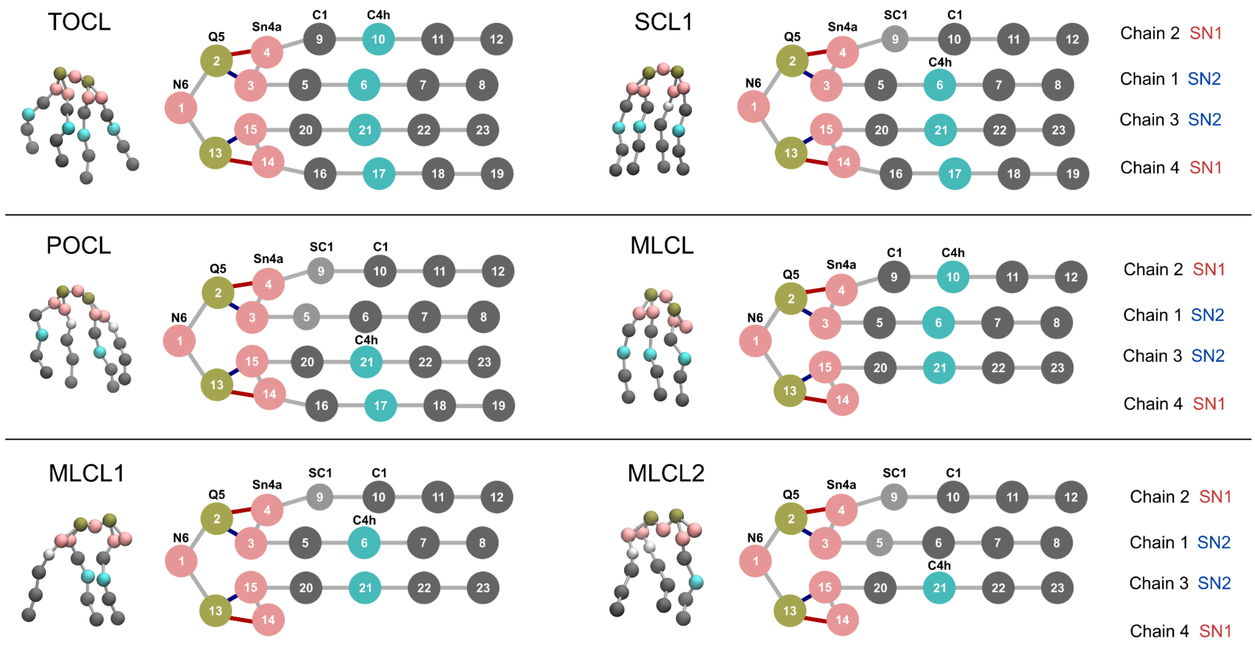


**Figure S3:** Generation of modified CL molecules in Martini3. Coarse-grained models for cardiolipin variants with Martini3 mapping. Head-group beads, identical in all cases, are shown in pink and tan. Each acyl chain is represented by beads according to chemical type: pink (carbonyl, ~2 atoms), light gray (first saturated segment, ~2 atoms), dark gray (saturated, ~4 atoms each), and cyan (unsaturated, ~4 atoms each). The parameters (bead types and bond types) to construct all molecules were taken from previously reported TOCL and palmitic acid values (1).

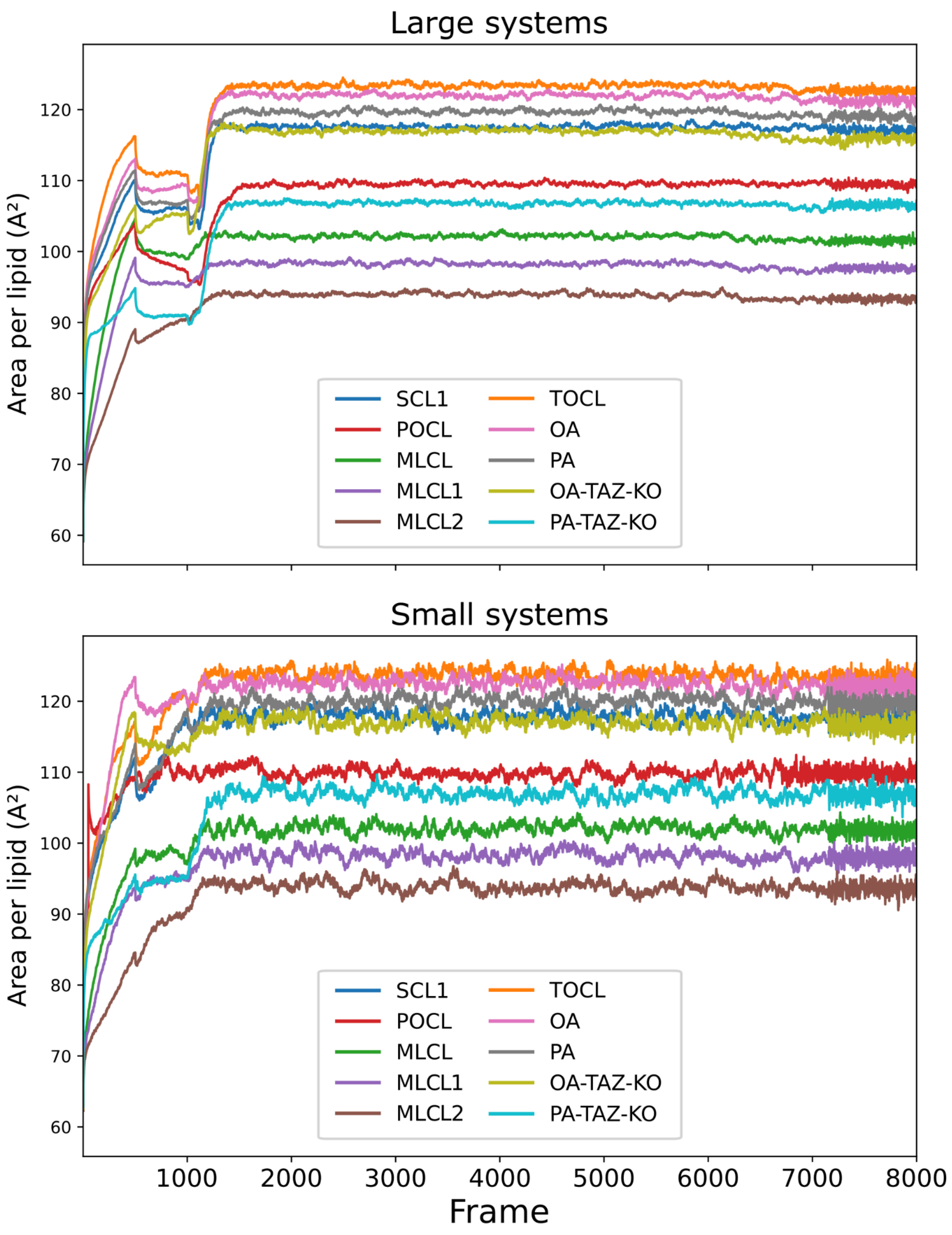


**Figure S4:** Area per lipid of modeled systems along the equilibration stages for large (top) and small (bottom) systems.

**Table S1:** Compositions of CG-MD systems. Values expressed as mol % of lipids.

| System | TOCL | SCL1 | POCL | MLCL | MLCL1 | MLCL2 |
|---|---|---|---|---|---|---|
| Pure TOCL | 100 | 0 | 0 | 0 | 0 | 0 |
| Pure SCL1 | 0 | 100 | 0 | 0 | 0 | 0 |
| Pure POCL | 0 | 0 | 100 | 0 | 0 | 0 |
| Pure MLCL | 0 | 0 | 0 | 100 | 0 | 0 |
| Pure MLCL1 | 0 | 0 | 0 | 0 | 100 | 0 |
| Pure MLCL2 | 0 | 0 | 0 | 0 | 0 | 100 |
| WT OA | 79 | 18 | 2 | 1 | 0 | 0 |
| WT Palm | 56 | 23 | 20 | 1 | 0 | 0 |
| TAZ KO OA | 51 | 25 | 4 | 15 | 5 | 0 |
| TAZ KO Palm | 4 | 13 | 53 | 1 | 20 | 9 |

**Table S2:** Number of lipid, water and ion beads for each CG-MD system.

| Large systems | | | | |
|---|---|---|---|---|
| System | Lipid beads | Water beads | $Na^+$ beads | Total |
| Pure TOCL | 124384 | 255553 | 10816 | 369054 |
| Pure SCL1 | 124384 | 255517 | 10816 | 390717 |
| Pure POCL | 124384 | 255510 | 10816 | 390710 |
| Pure MLCL | 102752 | 255486 | 10816 | 369054 |
| Pure MLCL1 | 102752 | 255482 | 10816 | 369050 |
| Pure MLCL2 | 102752 | 255471 | 10816 | 369039 |
| WT OA | 124122 | 255536 | 10812 | 390470 |
| WT Palm | 124076 | 255493 | 10808 | 390377 |
| TAZ KO OA | 120018 | 255550 | 10812 | 386380 |
| TAZ KO Palm | 117812 | 255474 | 10808 | 384094 |
| Small systems | | | | |
| System | Lipid beads | Water beads | $Na^+$ beads | Total |
| Pure TOCL | 16606 | 37642 | 1444 | 55692 |
| Pure SCL1 | 16606 | 37647 | 1444 | 55697 |
| Pure POCL | 16606 | 37646 | 1444 | 55696 |
| Pure MLCL | 13718 | 37649 | 1444 | 52811 |
| Pure MLCL1 | 13718 | 37647 | 1444 | 52809 |
| Pure MLCL2 | 13718 | 37666 | 1444 | 52828 |
| WT OA | 16490 | 37645 | 1436 | 55571 |
| WT Palm | 16536 | 37648 | 1440 | 55624 |
| TAZ KO OA | 15984 | 37663 | 1440 | 55087 |
| TAZ KO Palm | 15612 | 37681 | 1432 | 54725 |

**Table S3:** Major software and versions used in this study

| Software | Version | Reference |
|---|---|---|
| Gromacs | 2025.2 | (2) |
| MDAnalysis | 2.10.0 | (3) |
| numpy | 2.4.0 | (4) |
| Scipy | 1.16.3 | (5) |
| matplotlib | 3.10.8 | (6) |
| membrane-spectral-analysis | 0.1.0 | (7) |
| insane | 1.2.0 | (8, 9) |
| VMD | 1.9.4a55 | (10) |
